# A trait or a state - how consistent are tail biting phenotypes in pigs?

**DOI:** 10.1101/2023.02.16.528837

**Authors:** Jen-Yun Chou, Rick B. D’Eath, Dale A. Sandercock, Keelin O’Driscoll

## Abstract

The physiological, psychological and behavioural traits of tail biting/bitten pigs have been widely studied, with most research focusing on identifying traits to predict tail biting phenotypes (biters, victims, or uninvolved ‘neutrals’). However, it is not clear if these traits persist once pigs are no longer involved in performing or receiving tail bites. This study investigated whether there was a difference in behavioural responses to a novel object test (NOT) between pigs which were tail biting performers (BITER), recipients (VICTIM), or not involved in the biting events (NEUTRAL). We then investigated whether these differences in responses were still evident six weeks later, when tail biting was less prevalent. We hypothesised that biters would exhibit more behaviours indicative of boldness, but also stress, than victims or neutrals, and that these differences would still be present later. A total of 120 undocked pigs (BITER, n = 48; VICTIM, N = 48; NEUTRAL, n = 24; 60 male, 60 female) were selected for testing. At the time of the first test (one week after pigs were moved into the finisher house at 12 weeks of age; T1), the prevalence of tail biting peaked. The same test was repeated six weeks later (T2) when biting had eased. Each pig was tested separately in a novel arena. After a minute of habituation, a brush head was introduced by dropping it down from above, at which point the 5 min test began. A saliva sample was taken immediately before the habituation (baseline) and after each test to evaluate cortisol concentration. Direct continuous behaviour observation was conducted. Overall, salivary cortisol concentrations were higher after than before the NOT (*P* < 0.001), although VICTIM pigs had a reduced elevation in cortisol after the test (*P* = 0.02) compared to BITER and NEUTRAL pigs in T2. Between phenotypes, baseline saliva cortisol concentrations were similar. BITER pigs approached the object quicker than VICTIM pigs (P = 0.01), but also had more high-pitched vocalisations (P < 0.01), but these differences observed in T1 were no longer present in T2. The results suggested that the NOT was sensitive to detect different levels of behavioural response; however, differences in BITER pigs’ behavioural responses were transient and did not persist once biting behaviour ceased. The long-term consequence of chronic stress caused by being tail bitten was manifested in VICTIM pigs’ blunted cortisol elevation six weeks later, after severe tail biting events.

## 1. Introduction

Tail biting is a damaging behaviour in pigs where a biter pig manipulates, chews and bites a recipient pig’s tail. It often occurs in outbreaks, which can begin suddenly, and involve severe injurious tail biting events spreading from pig to pig (Chou et al., 2019). Tail biting involves two subjects (the biter and the victim) and the characteristics of and the interaction between the two, will affect the development and consequences of tail biting (Brunberg et al., 2016; Sambraus, 1985; Valros, 2018). If tail biters get rewarding feedback from performing this behaviour, they may persist in biting and cause greater damage (Taylor et al., 2010). On the other hand, if tail-bitten pigs (victims) are unresponsive to being bitten, and do not take evasive or protective actions, e.g., clamping down their tails, fleeing or retaliating by fighting, the severity of tail biting behaviour may also worsen (Taylor et al., 2010). Schrøder-Petersen and Simonsen (2001) categorised the risk factors of tail biting into internal (related to the pigs themselves) and external (related to the environment and management practices) ones. Since then, many studies have attempted to characterise traits such as growth performance, health status, immune function, stress level, personality and behavioural profiles of biters and victims (Brunberg et al., 2016; Li et al., 2016; Munsterhjelm et al., 2013a, 2013b; Valros et al., 2015; Zonderland et al., 2011b).

Besides tail biters and victims, there is a third category of pig which are not involved in either action, even during severe tail biting episodes. These “neutral” pigs have been observed to engage in fewer social interactions even before a tail biting outbreak, compared to biters and recipients (Brunberg et al., 2013a; Munsterhjelm et al., 2013a, 2016). Some studies further found differences in behaviour and gene expression in neutral pigs compared to pigs that were in groups without tail biting, suggesting that understanding how these pigs stay “neutral” even during severe tail biting events could mean that it may be possible to breed pigs that naturally avoid tail biting in the future (Brunberg et al., 2013b). The different “role” a pig plays in the event of tail biting, though, is not always stable. Sometimes the same individual pig may be both a biter and a victim (Valros, 2018; Zonderland et al., 2011a). However, in most studies, a general distinction can be made between these tail biting phenotypes when tail biting was severe (i.e., tail biter, victim or neutral; Brunberg et al., 2013b; Munsterhjelm et al., 2013a, 2013b; Zonderland et al., 2011b; Zupan et al., 2012).

The pigs’ personality is considered an important internal factor that can play a role in their propensity to tail bite, be tail bitten or stay neutral (Ursinus et al., 2014; Valros, 2018). One aspect of an animal’s personality is their behavioural response to a stressful situation, and this can be tested by using a range of behaviour tests which have been effectively applied to farm animals (Finkemeier et al., 2018; Nielsen, 2020). The novel object test (NOT) is one such test. The test can be conducted in the home pen, where a novel object is presented to a group of pigs, or more commonly, it is conducted in a novel arena with an individual pig (Nielsen, 2020). The latter imposes both the stress of experiencing an unfamiliar environment while being socially isolated, as well as the sudden forced exposure to an unfamiliar object. This method has previously been used to assess levels of fearfulness, boldness and stress coping ability in pigs (Finkemeier et al., 2018; Haigh et al., 2020; Reimert et al., 2014; Ursinus et al., 2013). In relation to tail biting, Ursinus et al., (2014) found that tail biters appear more fearful in the NOT (less time near the object, more time staying alert); on the contrary, Zupan et al., (2012) found biters tended to be more explorative, bold and novelty-seeking. It is worth noting that the former conducted the test in a novel arena while the latter in the pigs’ home pen. Both studies investigated the responses of pigs to the NOT immediately following severe tail biting events.

We are not aware of any previous studies that have investigated whether differences in response to novel object tests between tail biting phenotypes persist longer term. Consistency over time could indicate that these responses are inherent components of the pig’s personality (a trait). Differences that are only present at the time of tail biting could indicate a more transient response (a state), induced by involvement in tail biting. This study compared the response to the NOT of tail biters, victims, and neutral pigs, both when tail biting was evident (12 weeks of age), and 6 weeks later when it had eased (18 weeks of age). The hypotheses tested were that these phenotypes would respond differently to the NOT, with biters displaying more bold and novelty-seeking behaviours as well as signs of stress than the other pig types, and that these differences in their response would be consistent across both tests.

## 2. Materials and methods

The experiment was conducted as part of a larger study (Chou et al., 2020) at the Pig Research Facility in Teagasc, Moorepark, Ireland. All procedures were approved by the Teagasc Animal Ethics Committee (TAEC124/2016).

### 2.1 Animals and housing

Animals (N = 120) for the current study were a subset of those included in a larger study as stated above. In the aforementioned study, 672 pigs (Landrace × Large White) were used to investigate the effect of increased dietary fibre and a single enrichment device on tail biting (Chou et al., 2020). This larger study employed a 2 × 2 × 2 factorial design with two types of enrichment device in the weaner stage (a wooden post or a rubber floor toy), two types of enrichment device in the finisher stage (the same device as in the weaner stage or the alternative), and two levels of dietary fibre (standard: weaner 3.7% and finisher 5.9% of crude fibre or high: weaner 5.3% and finisher 11.6% of crude fibre). The pigs were born over two batches (i.e. two replicates) three weeks apart. All piglets were teeth-clipped to minimise facial lesions and sows’ udder injuries but were not tail docked. Male pigs were not castrated. Pigs were weighed periodically to record growth parameters.

All pigs were randomly assigned to eight treatments in pens of 14 pigs (48 pens in total, 6 pens per treatment combination) at weaning (4 weeks of age). Between pens, pigs were balanced for weight, sex (seven male and seven female) and litter mates. Pigs were fed *ad libitum* with dry pelleted feed by a single-spaced wet-dry feeder. The size of weaner and finisher pens were 2.4 m × 2.6 m and 4.0 m × 2.4 m respectively with fully-slatted floors. Pigs were moved to the finisher house at 12 weeks of age without further mixing.

As part of the study, direct behaviour observation on each pen of pigs was conducted on two different days every two weeks, starting from 7 weeks of age, and tail lesion scores were also recorded fortnightly for individual pigs from 8 weeks of age. A tail biting outbreak control protocol was employed (Chou et al., 2019), so all pens were checked multiple times daily and any tail biting event recorded in detail (i.e. biters, bitten pigs, treatments provided, etc.).

### 2.2 Pig selection for the NOT

Pigs were selected for the NOT at the time of moving into the finisher house (12 weeks of age). The timing of selection was based on suggestions from previous studies that the majority of tail biting events took place between 8-12 weeks of age (Haigh et al., 2019; Lahrmann et al., 2017; O’Driscoll et al., 2013; Schrøder-Petersen and Simonsen, 2001). Nineteen severe tail biting events were indeed recorded during this time (Chou et al., 2019). From each experimental pen (n = 48), one tail biter (BITER; n = 48), one tail bitten victim (VICTIM; n = 48), and one neutral pig (NEUTRAL; n = 24, pigs which were not involved in tail biting either as biters or victims) were selected.

The selection of test pigs was mainly based on their involvement in tail biting outbreaks, daily check-ups and behavioural observation records. In this study, we utilised a strict definition of ‘tail-biting outbreak’ with the criteria involving the highest level of tail damage. The tail damage scoring method was described in detail in Chou et al., (2019) and involved a 0 to 3 scale for tail damage, presence of blood and severity of tail amputation, with 0 being no damage, blood or amputation. Even without meeting the criteria to be considered an outbreak, most pens experienced enough biting behaviour that it was possible to identify at least one BITER and VICTIM pig.

A pig was categorised as BITER if it was a) identified as an active tail biter during a tail biting outbreak event, b) recorded as an active tail biter during routine check-ups the most frequently if no outbreak occurred in the pen, or c) noted as being the most frequent tail biter in the pen during the behaviour observation sessions conducted as part of the main study.

A pig was categorised as VICTIM if it a) was identified as a tail bitten victim during a tail biting outbreak event, b) required any medical treatment for tail wound if no outbreak occurred in the pen, c) recorded as having visible tail injuries during routine check-ups, or d) received the highest (i.e. worst) tail lesion score in the pen during the last tail scoring session (5 days before the test), if no pig satisfied any of the above criteria.

A NEUTRAL pig was selected if it was not identified as either BITER or VICTIM and had the lowest tail lesion score during the last tail scoring session (5 days before the test) in a pen. All selection criteria used records from between 8-12 weeks when most severe tail biting events took place. The sex of pigs in each category was balanced so that half of the tested pigs were male, and half were female. NEUTRAL pigs were not selected from the first replicate because of limitations in personnel, and inexperience in handling tail biting. A total of 120 pigs across the two replicates were selected (48 pigs in rep 1 and 72 pigs in rep 2). The sex, growth parameters and tail lesion scores of the selected phenotypes are detailed in **Table 1**.

**Table 1.** Information on the selected test pigs based on phenotypes. Post-hoc comparisons between phenotypes (LMM) after the Tukey-Kramer adjustment were indicated by different small letters. Tail lesion score was the average of the tail damage score (0-3 scale, 0: no lesion to 3: swollen bite wounds) and the tail blood score (0-3 scale, 0: no blood to 3: fresh blood). ADG: Average daily gain.

|  | BITER | NEUTRAL | VICTIM | *F*-value | *P*-value |
| --- | --- | --- | --- | --- | --- |
| Number of pigs (n) | 48 | 24 | 48 |  |  |
| Female | 24 | 12 | 24 |  |  |
| Male | 24 | 12 | 24 |  |  |
| Growth (kg) |  |  |  |  |  |
| Weaning weight | 7.71 ± 0.20 | 7.96 ± 0.30 | 7.82 ± 0.20 | 0.26<sub>(70.3,2)</sub> | NS |
| Moving weight | 37.00 ± 0.86 | 38.47 ± 1.30 | 38.19 ± 0.86 | 0.76<sub>(66.7,2)</sub> | NS |
| Finishing weight | 96.54 ± 1.67 | 96.20 ± 2.57 | 97.32 ± 1.75 | 0.09<sub>(67.7,2)</sub> | NS |
| Weaner ADG | 0.60 ± 0.02 | 0.62 ± 0.02 | 0.62 ± 0.02 | 0.75<sub>(66.4,2)</sub> | NS |
| Finisher ADG | 1.08 ± 0.02 | 1.05 ± 0.03 | 1.07 ± 0.02 | 0.40<sub>(65.9,2)</sub> | NS |
| Overall ADG | 0.85 ± 0.01 | 0.85 ± 0.02 | 0.86 ± 0.02 | 0.12<sub>(67.4,2)</sub> | NS |
| Tail lesion score (0-3) |  |  |  |  |  |
| Week 12* | 0.49 ± 0.09<sup>a</sup> | 0.35 ± 0.12<sup>a</sup> | 1.75 ± 0.09<sup>b</sup> | 73.27<sub>(113,2)</sub> | < 0.001 |
| Week 18† | 0.55 ± 0.05 | 0.43 ± 0.08 | 0.67 ± 0.06 | 2.86<sub>(68,2)</sub> | 0.06 |
| Weaner stage average | 0.59 ± 0.05<sup>a</sup> | 0.62 ± 0.07<sup>a</sup> | 1.40 ± 0.05<sup>b</sup> | 100.1<sub>(72.4,2)</sub> | < 0.001 |
| Finisher stage average | 0.53 ± 0.06<sup>a</sup> | 0.52 ± 0.08<sup>a</sup> | 0.78 ± 0.06<sup>b</sup> | 7.35<sub>(77.8,2)</sub> | < 0.01 |

\*Week 12 is the time when pigs were moved into the finisher house and also when the tail biting phenotypes were selected and tested in the first test (T1).
†Lesion scores were taken just before the second test (T2) in week 18.

### 2.3 NOT procedures

The NOT procedure used in the current study was adapted from Haigh et al. (2020). The test was conducted twice on each test pig. The first test was conducted at the end of 12 weeks of age, one week after pigs were moved into the finisher house (T1), during the period when tail biting was most frequent. The second test was conducted six weeks later (T2) at 18 weeks of age (one week before slaughter) when severe tail biting had ceased. Seven test pigs did not proceed to T2 due to being permanently removed from the main study as a result of earlier severe tail biting (6 victims, 2 female/4 male, and 1 male obsessive biter; T1: n = 48, T2: n = 41). As there were two batches of pigs born 3 weeks apart in the main study, this NOT testing also took place in two phases 3 weeks apart.

The testing took place over three consecutive days between 1300-1800h. The test pen was located in an isolated, empty and unused room inside the same finisher building, without audio, visual or olfactory contact with other pigs. The floor of the test pen (4.0 m × 2.4 m) was fully slatted and divided into nine squares by spray paint on the floor to record pigs’ locomotion inside the pen. The novel object used was a wooden sweeping broom head which had bright orange PVC bristles (0.32 × 0.15 × 0.09 m) (Haigh et al., 2020). The introduction of the broom head into the pen was done using a pulley system on the ceiling and a PVC cord. This was to facilitate controlled lowering of the object into the pen by the experimenter who was located outside the pen, without additional disturbance to the test pig (**Figure 1**). The same novel object was used in both T1 and T2. The order of testing different pens and categories of pigs within a pen was randomised.

**Figure 1.**
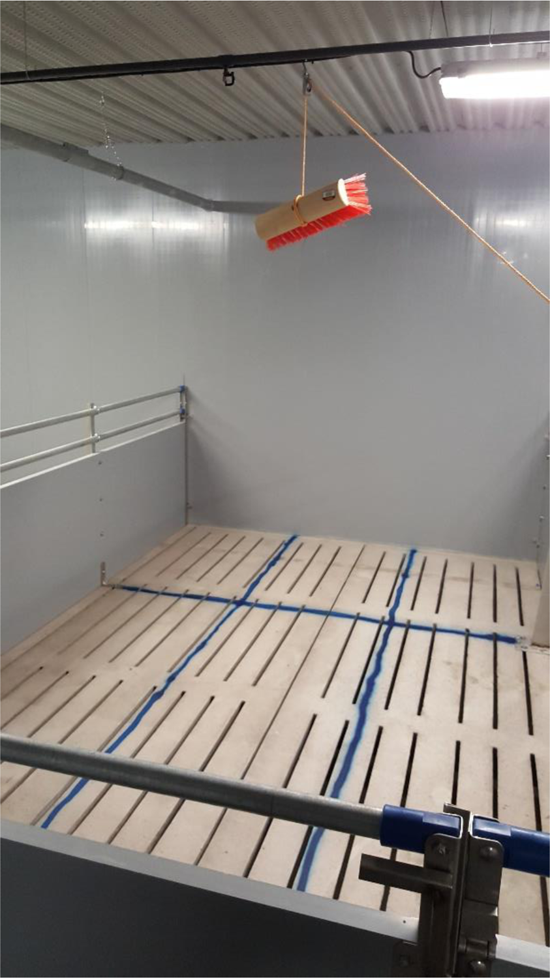
Photograph of the novel object test (NOT) arena, the novel object (broom head) and pulley system to facilitate the introduction and removal of the novel object into the test arena.

All test pigs in the pens were separated from their pen mates calmly and along with a companion pig which voluntarily moved out of the pen with the test pigs, then brought to a waiting area. This was done so that during waiting the test pig was not at the risk of solitary confinement before the test since there was always at least one pig as a companion. Immediately prior to the test, a saliva sample was obtained from the test pig using a biocompatible synthetic swab (Salivette, Sarstedt, Wexford, Ireland) presented by metal tweezers for chewing. As the pigs were familiar with the experimenter, chewing the swab could be done voluntarily without inducing extra stress to the pig, taking around 30 seconds. The test pig was then guided into the test room and acclimatised to the test pen for 1 minute before the novel object was lowered to the centre of the pen and the behaviour recording began. Observation continued for a further 5 minutes, and afterwards a second saliva sample was obtained from the pig inside the test pen before it was led to leave the room and returned to its home pen. Any urine or faeces were cleaned by hosing down the slats prior to the next test.

### 2.4 Measurements

#### 2.4.1 Salivary sampling

The saliva samples obtained before and after the test (volume of approximately 0.5 ml) were preserved in storage tubes (Salivette, Sarstedt, Wexford, Ireland), centrifuged at 1,500 rpm and then frozen at −20ºC. The samples were analysed using ELISA (Enzyme-linked immunosorbent assay, Salimetrics, Carlsbad, CA, USA; 96-well plate with assay sensitivity of 0.007 μg/dL and assay range between 0.012-3.000 μg/dL) to quantify the cortisol concentration in the saliva (25 μL of samples/test) (Salimetrics, 2021; Thomsson et al., 2014). The average coefficient of variation (CV) calculated from the means of high and low plate control samples (inter-assay) was 24.6% and the average CV between duplicates within plates (intra-assay) was 3.6%.

#### 2.4.2 Behaviour recordings

Behaviour of the test pig was recorded continuously throughout the 5-min test, which commenced after one minute of habituation (a total test time of six minute). Behaviours recorded consisted of state events and point events, all coupled with location coding by the same human observer using the Psion Workabout Pro 4 (Motorola, Chicago, USA) with the Observer software (Noldus Information Technology, Wageningen, The Netherlands). State events were mutually exclusive and recorded as durations, whereas point events were recorded as frequencies and could occur simultaneously with state events (**Table 2**).

**Table 2.** Ethogram for behaviour observation in the NOT. State events were recorded as durations of time and frequencies, and point events as occurrences.

| Behaviour | Description |
| --- | --- |
| **State event** |  |
| Interaction with the object | Being in contact with the object with mouth, snout or any other body part |
| Attention* | Standing still with attention towards the object without contact with it |
| Withdraw | Displaying backwardly reactive movement due to the object |
| Stationary* | Standing still |
| Sitting/lying | Sitting with either all fours legs down (lying) or with its two hind legs down (sitting) |
| Walking<sup>†</sup> | Moving around the pen with head raised |
| Exploring floor<sup>†</sup> | Moving around the perimeter of the pen, exploring the ground with its snout or mouth (direct contact or close proximity) |
| Exploring wall<sup>†</sup> | Moving around the perimeter of the pen, exploring the wall with its snout or mouth (direct contact or close proximity) |
| Play | Scampering, pivoting, head tossing, flopping, and pawing movement |
| **Point event** |  |
| Low-pitch vocalisation | Vocalisation with a low pitch tone (grunt) |
| High-pitch vocalisation | Vocalisation with high pitch tone (squeal or grunt squeal) |
| Elimination | Defecating or urinating |
| Jump | Jumping in air or against a wall of the arena |

\* Stationary and attention were combined into staying alert during statistical analysis.
<sup>†</sup> Walking, exploring floor and exploring wall were also combined as a new category “exploration” and analysed.

### 2.5 Data analysis

Data were analysed using Statistical Analyses System (SAS, version 9.1.3, 1989, SAS Institute Inc., Cary, NC, USA). Linear mixed models (LMM) were used to analyse the data unless specified otherwise. The Tukey-Kramer adjustment was used to detect post-hoc differences between least squared means.

Comparisons between phenotypes for body weights, average daily gains and tail lesion scores at different ages were conducted using LMM, including all treatments in the main study, replicate, phenotype (BITER, NEUTRAL or VICTIM) and all interactions as fixed effects, with pen as a random effect.

For analysis of salivary cortisol: Residuals were right-skewed, so the data were square root transformed before analysis to improve normality. All treatments in the main study, replicate, phenotype, test (T1 or T2), time of collection (before or after the test) and all interactions were included in the model (LMM) as fixed effects. Time of collection nested within test was the repeated effect, and pen and the plate on which the ELISA was analysed were also included as random effects. Step-wise removal of explanatory variables was carried out until the best-fit model was confirmed by the smallest Akaike’s Information Criterion (AIC) (Ngo and Brand, 1997). The contrast function was used to compare each phenotype with the other two types.

For analysis of the behavioural data: The model (LMM) included all treatments in the main study, replicate, phenotype, test and all interactions as fixed effects, test as the repeated effect and pen as the random effect. Walking, exploring floor and exploring wall were further combined as “exploration” and analysed, as were stationary and attention as “staying alert.” These are based on a previous study (Haigh et al., 2020) and biological meaningfulness. The latency to approach the novel object and the number of occurrences of high-pitch vocalisation were transformed by logarithm to improve normality. The same step-wise removal described above was used to find the best-fit model. The contrast function was used to compare one phenotype against the other two.

The likelihood to display a withdraw reaction from the object between tests was analysed using generalised linear mixed model with a binary distribution (with/without withdraw reaction) and the logit link function to check the odds ratio. Test was included as the repeated effect and pen as the random effect.

Data from T1 and T2 were also analysed independently to understand any test-specific differences and whether any differences detected between phenotypes were consistent across both tests.

## 3. Results

The comparisons between phenotypes showed that there was no difference in any growth parameter (**Table 1**). As expected, and in line with the study design, tail lesion scores were higher for VICTIM when T1 was conducted but were not different from BITER or NEUTRAL in week 18, when T2 was conducted (**Table 1**).

### 3.1 Salivary cortisol

Overall, pigs’ saliva cortisol concentrations were lower before being tested in the NOT than after (0.20 ± 0.03 vs. 0.33 ± 0.03 μg/dL, *P* < 0.001) across all phenotypes. Overall values were higher in T2 than in T1 (0.31 ± 0.03 vs. 0.21 ± 0.03 μg/dL, *P* < 0.001). The magnitude of the increase in salivary cortisol concentration before to after the NOT did not differ between T1 and T2.

There was no difference in baseline salivary cortisol concentrations (before NOT) between the selected phenotypes. However VICTIM pigs had a lower elevation in salivary cortisol concentrations after the NOT compared to the other two types of pigs combined (0.09 ± 0.03 μg/dL vs. BITER 0.15 ± 0.03 μg/dL and NEUTRAL 0.18 ± 0.04 μg/dL, *F*_(69,1)_ = 5.35, *P* = 0.02). When considering T1 and T2 separately, this effect was only present in T2 (VICTIM 0.11 ± 0.04 μg/dL vs. BITER 0.23 ± 0.04 μg/dL and NEUTRAL 0.22 ± 0.05 μg/dL, *F*_(69,1)_ = 5.35, *P* < 0.01), but not in T1.

### 3.2 Behaviour

The most common state event observed was exploration (Walking + Exploring floor + Exploring wall), followed by being stationary, and then object interaction. Play behaviour was only observed being performed by one pig (VICTIM) and similarly, only three pigs were recorded sitting down (one BITER and two VICTIM). Twelve occurrences of jumping were recorded (two BITER, five NEUTRAL and five VICTIM pigs), and the number of occurrences was only numerically higher in NEUTRAL pigs (average 3.8 times compared to BITER 1 time and VICTIM 1.6 times).

#### 3.2.1 Differences between phenotypes

Overall, BITER pigs approached the object quicker than VICTIM pigs (**Figure 2**, *P* = 0.01) and also performed more high-pitch vocalisations compared to the other two types of pigs (**Figure 2**, *P* < 0.01).

**Figure 2.**
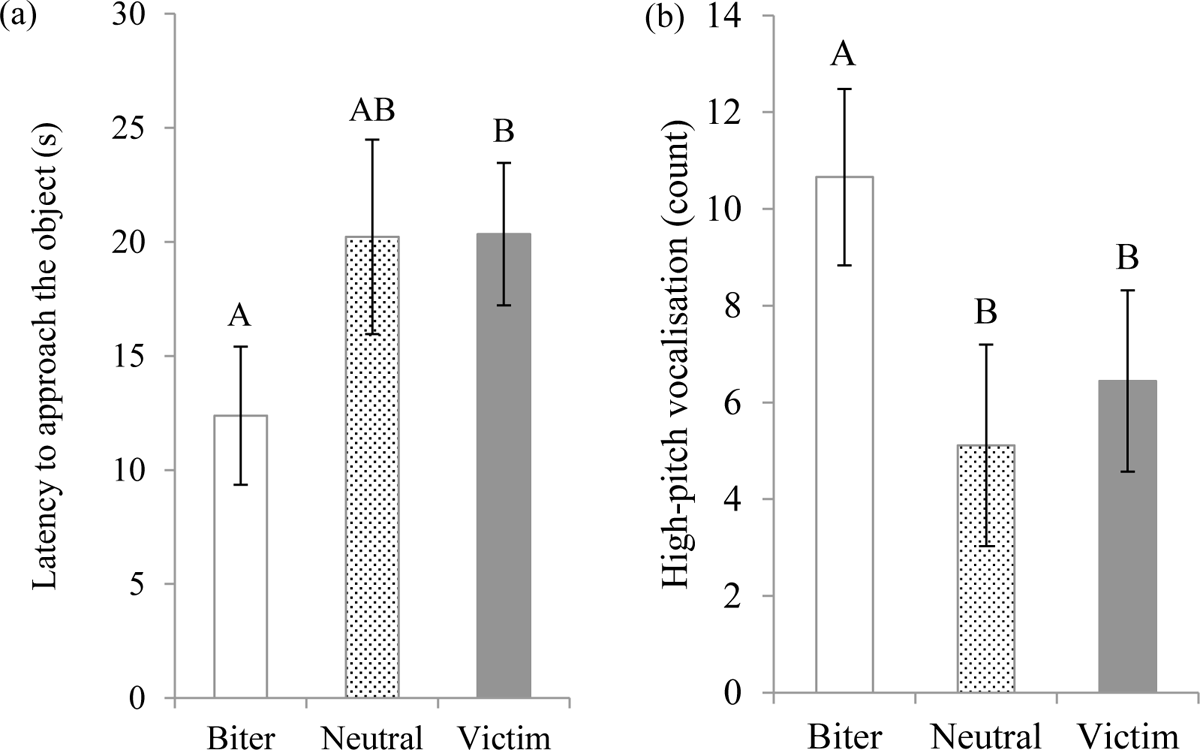
Comparisons between the three tail biting phenotypes in behaviours across both tests combined. 3(a): time (s) taken to approach the novel object after the NOT began (*F*_(78,2)_ = 4.46, *P* = 0.01) and 3(b): the number of occurrences of high-pitch vocalisation (count) (*F*_(129,1)_ = 8.09, *P* < 0.01). Post-hoc comparisons after the Tukey-Kramer adjustment were indicated by different capital letters. Data are presented as LSMeans ± S.E. (*n* = 120).

When analysing the data from T1 and T2 separately, BITER pigs approached the object faster in T1 than VICTIM pigs (**Figure 3**, *P* = 0.01), but there was no difference in T2 anymore. BITER pigs also performed more frequent high-pitch vocalisations in T1 compared to the other two phenotypes combined (**Figure 3**, *P* = 0.05), but this difference was no longer significant in T2 either.

**Figure 3.**
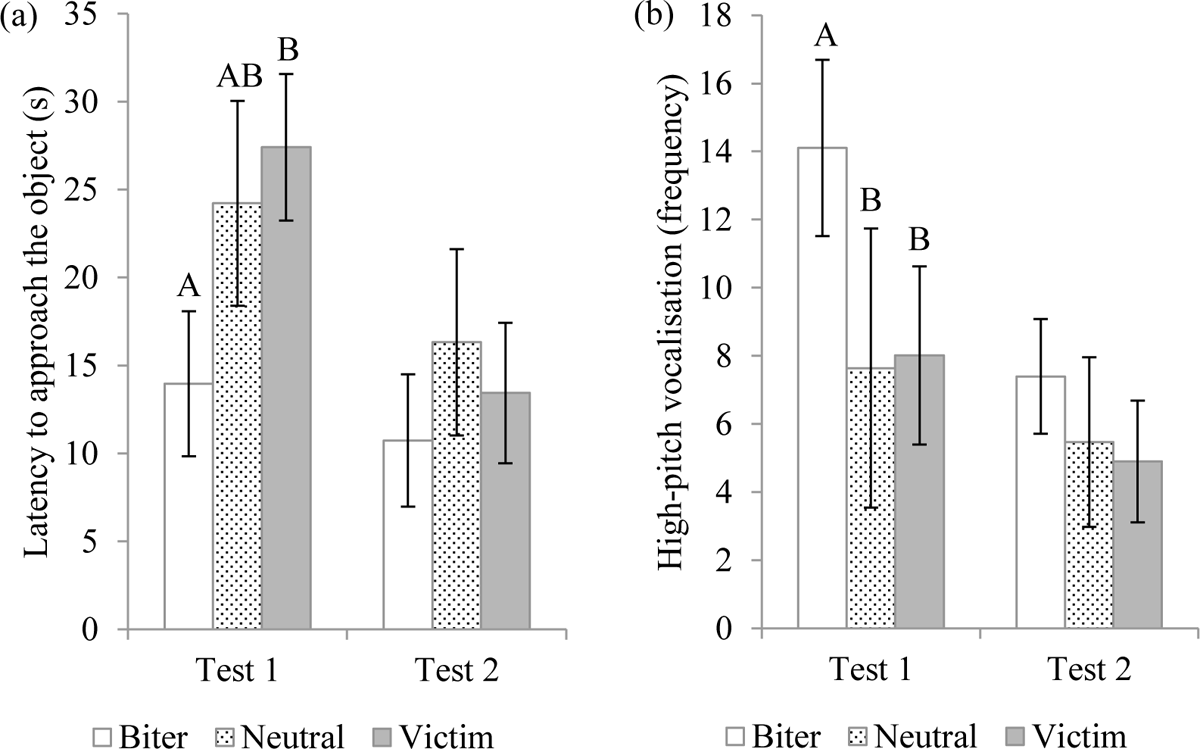
Comparisons between the three tail biting phenotypes in T1 (12 weeks) and T2 (18 weeks): a) Time (s) taken to approach the novel object after its introduction into the test arena (T1: *F*_(78.1,2)_ = 4.59, *P* = 0.01; T2: NS) and b) Number of high-pitch vocalisations (T1: *F*_(107,1)_ = 3.83, *P* = 0.05; T2: NS). Post-hoc comparisons after the Tukey-Kramer adjustment were indicated by different capital letters. Data are presented as LSMeans ± S.E. (*n* = 120).

#### 3.2.2 Differences between tests

Pigs on average performed more high-pitch vocalisations in T1 than in T2 (**Table 3**, *P* < 0.001). They were also 2.17 times more likely to display a withdraw reaction from the object in T1 compared to T2 (OR estimate = 8.76, 95% CI 4.10-18.72, *P* < 0.001). In contrast, they spent less time staying alert (Stationary + Attention, *P* < 0.01) and also approached the novel object sooner in T2 than T1 (*P* < 0.001, **Table 3**). However, they also spent more time near the gate side of the test arena in T2 (*P* < 0.001, **Table 3**). There were no other differences in behaviour between the two test days.

**Table 3.** Behavioural comparison between T1 (conducted when pigs were 12 weeks of age) and T2 (conducted when pigs were 18 weeks of age). Latency to approach the novel object and the number of occurrences of high-pitch vocalisation were log-transformed for the analysis so the back-transformed LSMeans ± S.E. were displayed. Staying alert is a combination of being stationary and standing still paying attention to the object without interacting with it.

| Behaviour | T1 | T2 | *F*-value | *P*-value |
| --- | --- | --- | --- | --- |
| Object interaction (s) | 83.61 ± 4.60 | 75.58 ± 4.69 | 2.00<sub>(112,1)</sub> | NS |
| Latency to approach the object (s) | 21.93 ± 2.76 | 13.36 ± 2.81 | 23.73<sub>(114,1)</sub> | < 0.001 |
| Staying alert (s) | 101.40 ± 4.67 | 82.10 ± 4.77 | 11.01<sub>(112,1)</sub> | < 0.01 |
| Near the gate of the arena (s) | 63.17 ± 4.18 | 93.57 ± 4.26 | 33.59<sub>(114,1)</sub> | < 0.001 |
| High-pitch vocalisation (counts) | 11.01 ± 1.47 | 5.12 ± 1.49 | 17.31<sub>(112,1)</sub> | < 0.001 |

#### 3.2.3 Differences between treatments in the main study

The treatment in the main study had an effect on the amount of time interacting with the object in both T1 and T2. Pigs with the rubber floor toy in the weaner stage interacted with the novel object for longer in T1 (97.09 ± 6.28 vs. the spruce post 74.27 ± 6.22 seconds, *F*_(107,1)_ = 7.5, *P* < 0.01) and those with the rubber floor toy in the finisher stage also interacted for longer with the object in T2 (83.35 ± 6.65 vs. the spruce post 62.66 ± 6.88 seconds, *F*_(34,1)_ = 5.06, *P* = 0.03).

## 4. Discussion

This study investigated the responses of three pig tail biting phenotypes to a novel object test (NOT) at 12 weeks of age and whether any difference in response was consistent in a second test at 18 weeks of age after the performance of and effects of tail biting were diminished. Similar to the initial hypothesis, there were differences in pigs’ responses both in terms of saliva cortisol concentration and behaviour; VICTIM pigs exhibited a blunted elevation in salivary cortisol concentration after the test compared to BITERS and NEUTRALs relative to pre-test baseline, and BITER pigs approached the novel object quicker but also performing more high-pitched vocalisations. However, BITER pigs’ behavioural differences were only evident in T1 and not anymore in T2, which suggests that the responses identified in T1 were likely due to the immediate consequence of ongoing or recent tail biting. On the other hand, the VICTIM pigs’ blunted increase in cortisol reacting to NOT was more pronounced in T2, suggesting a longer-term effect of tail biting. The lack of consistency across time indicates that these responses are likely due to temporary or imposed physical or psychological states, rather than stable traits.

The overall increase in saliva cortisol concentration after the NOT was expected as the test is designed to impose an elevated state of arousal. Social isolation in an unfamiliar arena without audio, visual and olfactory contact with conspecifics can elicit a behavioural and physiological fear response in pigs (Forkman et al., 2007; Murphy et al., 2014). This effect can be augmented when a novel object is presented to them in an unexpected fashion (i.e. dropped from above after being in the novel arena for 1 minute), and an increase in salivary cortisol levels after an open field test or NOT has been shown in previous studies (Haigh et al., 2020; Reimert et al., 2014). For prey species, suddenness and unpredictability are key elements in predation and therefore can stimulate innate fear and stress responses (Forkman et al., 2007). However, another explanation is that salivary cortisol concentration was elevated, in part, due to mental or physical arousal during the test, rather than fear *per se*. Pigs spent less time standing still in T2, and this could be the reason for the higher cortisol concentration recorded in T2 compared to T1. Either way, the observed elevations in saliva cortisol concentrations in response to the NOT were within a similar range to those reported by (Thomsson et al., 2014) for primiparous gilts and multiparous sows subjected to group mixing for 12 hours (median 0.66; range 0.10-1/17 ug/dL and median 0.47; range 0.14-2.53 ug/dL respectively). This model for social stress has been shown to activate the hypothalamo-pituitary-adrenal (HPA) axis in pigs (Rutherford et al., 2009; Soede et al., 2006), and therefore it is reasonable to assume that the HPA axis was activated to some extent by the NOT in the pigs investigated in current study.

Between tail biting phenotypes, there was no difference in the baseline saliva cortisol concentration at either test, but the overall magnitude of increase in cortisol concentration after the test was lower in VICTIM pigs compared to the other two pig types combined. Although it may seem that these bitten pigs were less stressed in the NOT, chronically stressed animals can have a lower cortisol response to an acute stressor (Janssens et al., 1995). In rats, maternal deprivation led to blunted ACTH levels after a stressful event later in life and could also increase anxiety-life behaviours (Daniels et al., 2004). Tail bitten pigs have been shown to suffer from chronic stress with altered physiological responses (Munsterhjelm et al., 2013a; Valros et al., 2015, 2013). This may explain why the difference in cortisol elevation was only different in T2 as it takes time to develop the symptoms of chronic stress. It is interesting that VICTIM pigs were less responsive to the test in T2, even though their tail lesion scores were no longer different to those of BITERS and NEUTRALS and were mostly healed up. Thus, even after physical healing occurs, it is possible that the experience of being badly bitten could lead to less easily observable long term effects for the pig, whether due to psychological or physical stress.

On the other hand, similar to what Zupan et al., (2012) reported, BITER pigs approached the novel object more quickly than VICTIMs which is an indication of boldness and novelty seeking. However, they also squealed more frequently than the other two types of pigs. As mentioned previously, high-pitch vocalisations could be seen as signs of stress under a challenging situation. Some suggested animals with a proactive coping style can display bolder behaviours but may adapt poorly to a new environment (Kanittz et al., 2019). This also agrees with previous studies that tail biters may experience stress which manifests in damaging behaviours (Munsterhjelm et al., 2013a; Ursinus et al., 2014; Valros et al., 2013). The NEUTRAL pigs did not show any pattern of response to the test that was distinguishable from the other two phenotypes.

Despite the fact that there were overall differences in behaviours between BITER pigs and the others, when the data from T1 and T2 were analysed independently, no difference in behaviours between pig phenotypes was present in T2. This suggested that any impact of pigs’ involvement in tail biting as a BITER that was reflected in response to the NOT was only transient, and occurred only when the level of tail biting was severe. T1 was conducted when severe tail biting was ongoing, and the behavioural differences during the test might reflect their ‘state’ of mind when they were engaged in tail biting as per their role in tail biting (BITER, VICTIM, or NEUTRAL). The fact that these differences were no longer present 6 weeks later suggests that the tail biting phenotype is not an underlying, stable trait for pigs as hypothesised. This lack of a stable individual difference in pigs’ propensity to perform tail biting, is different to other damaging behaviours, such as the performance of aggression in pigs, which has been shown to exhibit a high consistency and stability over time (Clark and D’Eath, 2013). It is worth noting that pigs’ involvement in tail biting can sometimes be fluid and change over their lifetimes (Ursinus et al., 2014; Zonderland et al., 2008), which may also explain the lack of consistency in the response to the test. In addition, as seven most extreme BITER and VICTIM pigs were removed prematurely from the study due to humane endpoints and did not participate in T2, this could have also influenced the outcome of T2.

Pigs spent less time in an alert state during T2 than T1. Donald et al., (2011) found that when pigs were injected with an anti-stress agent, they became more active in an open field test, which suggested that being less stationary in the open field is a sign of less of a stress response. High-pitch vocalisations were also recorded more frequently in T1, and these are usually associated with heightened adrenaline, and an aversive response (Düpjan et al., 2008; Fraser, 1974; Reimert et al., 2013). In terms of object interaction, pigs approached the object faster in T2 and were also less likely to withdraw from it. These data indicate that pigs showed some level of habituation to the test in T2. In order to reduce habituation due to repeated testing, some suggested maintaining the stimulation of the novel object by using different objects (Forkman et al., 2007). In the current study, the same object was used in both T1 and T2 to avoid any confounding effect that could be brought about by the choice of the object itself. This might have decreased the novelty of the “novel object.” However, previous work has suggested that pigs’ memory retention on familiar object can last for one hour (Kornum et al., 2007) or up to 48 hours (Fleming et al., 2018), and other research found pigs retained no memory of an object if the object was only exposed previously for 10 minutes (Gifford et al., 2007). No evidence suggested that pigs can recognise the same object after an interval of 6 weeks with only a 5-minute exposure previously. Therefore, it was considered suitable to reuse the same object for both tests.

There was an effect of the enrichment treatment from the main study (Chou et al., 2020). Overall, when pigs were provided with the rubber floor toy as environmental enrichment during the period before the test took place, they spent more time interacting with the novel object. This may be because the presentation of the novel object resembles that of the rubber floor toy, which was placed on the floor of the pen and movable. The familiarity with floor enrichment items that can be moved around could lead the pigs to interact with the novel object more frequently. Thus factors such as the prior experience of the pigs should be considered carefully when planning novel object tests.

## 5. Conclusions

This study found that the three pig phenotypes most commonly associated with tail biting (BITER, NEUTRAL and VICTIM) reacted differently to a NOT in terms of elevation of salivary cortisol concentration and behaviour. The behavioural differences between BITER pigs and the other phenotypes no longer existed in T2, when the scale of tail biting was less severe, and on the other hand, VICTIM pigs exhibited blunted reaction in cortisol elevation only in T2, when tails were healed. Thus the effects on an individual pig of being badly bitten may induce chronic stress and last much longer than the physical healing of the tail. Overall, this study suggests traits associated with the tail biting phenotypes were transient and not consistent over 6 weeks when levels of tail biting severity differed.

## Acknowledgements

This project was co-funded by the Teagasc Walsh Fellowship, the Department of Agriculture, Food and the Marine in Ireland. SRUC also receives funding from the Rural & Environmental Science & Analytical Services Division of the Scottish Government. We would like to acknowledge the help during the tests from intern students Alexandra Courty and Marleen van de Heide.

## Data, scripts, code, and supplementary information availability

Data and statistical scripts used in this study is deposited in the OSF depository and available at https://osf.io/qh6ze/.

